# p.D372H: A novel SCN5A mutation associated with Brugada syndrome

**DOI:** 10.1101/2025.01.29.635599

**Authors:** Xianghuan Xie, Yanghui Chen, Zhiqiang Li, Xian-Cheng Jiang, Yang Sun, Guangzhi Chen

**Author notes:** Corresponding authors Guangzhi Chen, Division of Cardiology, Department of Internal Medicine, Tongji Hospital, Tongji Medical College, Huazhong University of Science and Technology, 1095# Jiefang Ave., 430030, Wuhan, China; Hubei Key Laboratory of Genetics and Molecular Mechanisms of Cardiological Disorders, Wuhan 430030, China., Yang Sun, Division of Cardiology, Department of Internal Medicine, Tongji Hospital, Tongji Medical College, Huazhong University of Science and Technology, 1095# Jiefang Ave., 430030, Wuhan, China; Hubei Key Laboratory of Genetics and Molecular Mechanisms of Cardiological Disorders, Wuhan 430030, China.

## Abstract

**Background:** Brugada syndrome (BrS) is a genetic cardiac arrhythmia disorder inherited in an autosomal dominant manner, characterized by ST-segment elevation in the right precordial leads (V1-V3) on electrocardiograms (ECGs). This syndrome predominantly affects young individuals with structurally normal hearts and significantly increases the risk of ventricular arrhythmias and sudden cardiac death (SCD). The most common genotype found among BrS patients is caused by mutations in the SCN5A gene, which lead to a loss of function of the cardiac sodium (Na+) channel (Nav1.5) by different mechanisms.

**Methods:** Plasmids containing SCN5A were constructed using PCR and site-directed mutagenesis to create the D372H mutation. HEK293 cells were cultured and transfected with the wild-type and mutant constructs. Patch-clamp recordings assessed sodium current characteristics. Confocal microscopy visualized channel localization. Quantitative RT-PCR analyzed mRNA expression levels, while Western blot evaluated protein expression using specific antibodies. We identified a novel missense mutation, D372H, in the SCN5A gene associated with Brugada syndrome. Functional assays in HEK293 cells expressing the D372H mutant revealed a near-complete loss of sodium currents. Subsequent experiments with co-transfection of WT and D372H plasmids demonstrated that co-expression led to a significant reduction in current density in WT-expressing cells (P < 0.05). The D372H mutation also resulted in a hyperpolarizing shift of approximately 20 mV in the voltage dependence of inactivation, while activation and recovery kinetics remained unaffected. Additionally, confocal microscopy showed reduced membrane localization of the D372H mutant, with a significant decrease in protein expression levels confirmed by Western blot and RT-qPCR analyses.

**Conclusion:** In summary, our findings indicate that the D372H mutation in the Nav1.5 sodium channel leads to significant reductions in sodium current density, altered channel expression, and impaired membrane localization. These changes contribute to the pathophysiology of Brugada syndrome by disrupting cardiac action potential dynamics.

## Introduction

Brugada syndrome (BrS) is a genetic cardiac arrhythmia syndrome inherited in an autosomal dominant manner, characterized by distinctive electrocardiogram (ECG) changes, specifically ST-segment elevation in the right precordial leads (V1-V3)^1^. This condition primarily affects young individuals with structurally normal hearts and is associated with an increased risk of ventricular arrhythmias and sudden cardiac death (SCD)^2,3^. BrS is estimated to account for 4-12% of all sudden death cases and approximately 20% of SCD occurrences in individuals under 50 years of age with normal cardiac structures^4,5^. The prevalence of BrS ranges from 1 in 2,000 to 1 in 5,000, with symptoms typically emerging in adulthood; the average age of SCD is reported to be 41 ± 15 years^6^.

To date, more than 500 variants have been discovered in over 40 genes that may be associated with Brugada syndrome (BrS)^7^. These variants primarily involve genes that encode sodium^8^, potassium^9^, and calcium^10^ channels, as well as related regulatory proteins. Nonetheless, the majority of these genetic alterations have not been validated. Currently, the Sodium Voltage-Gated Channel Alpha Subunit 5 (SCN5A)gene is the only gene definitively linked to BrS, responsible for less than 30% of cases, which leaves more than 60% of patients without an identified genetic cause^11,12^.

The voltage-dependent cardiac sodium channel, Nav1.5, plays a critical role in generating action potentials in cardiac muscle. Encoded by the SCN5A gene, Nav1.5 consists of α-subunit made up of 2,016 amino acids organized into four homologous domains (DI-DIV)^13,14^. Each of these domains contains six transmembrane α-helices (S1-S6), which are interconnected by intracellular loops. The voltage-sensing apparatus is primarily formed by the S1-S4 segments, which incorporate positively charged amino acids essential for channel activation. In contrast, the S5 and S6 segments, along with the S5-S6 loops, collectively create the channel’s pore and the selectivity filter for sodium ions (Na+). While SCN5A mutations linked to Brugada syndrome (BrS) are dispersed throughout the entire channel structure, a combination of structural and functional analyses has identified that certain mutations in specific regions are associated with distinct biophysical defects.

In our previous research, we discovered a novel mutation, NM_198056.2: c.372G>C (p.D372H)^15^, situated in domain I of the Nav1.5 protein, specifically within the extracellular loop that connects the S5 and S6 transmembrane segments (Figure 1)^16,17^. In 2020, Tadashi Nakajima and colleagues identified the SCN5A W374G mutation, which is located in the pore loop of Domain I, in cases of severe Brugada syndrome (BrS) and is in close proximity to the D372H mutation. Their functional studies indicated that the W374G mutation led to a reduced current density, likely due to trafficking defects, and a depolarizing shift in steady-state activation (SSA), highlighting the biophysical abnormalities associated with mutations in this region^18^. These findings suggest that mutations within this domain significantly disrupt Na^+^ channel function; however, the impact of the D372H mutation on channel activity has not been thoroughly investigated. The objective of this study is to clarify the relationship between the D372H mutation and Brugada syndrome. Through comprehensive genetic analyses and detailed electrophysiological assessments, we aim to deepen our understanding of how novel SCN5A mutations contribute to the pathophysiology of Brugada syndrome.

**Fig 1.**
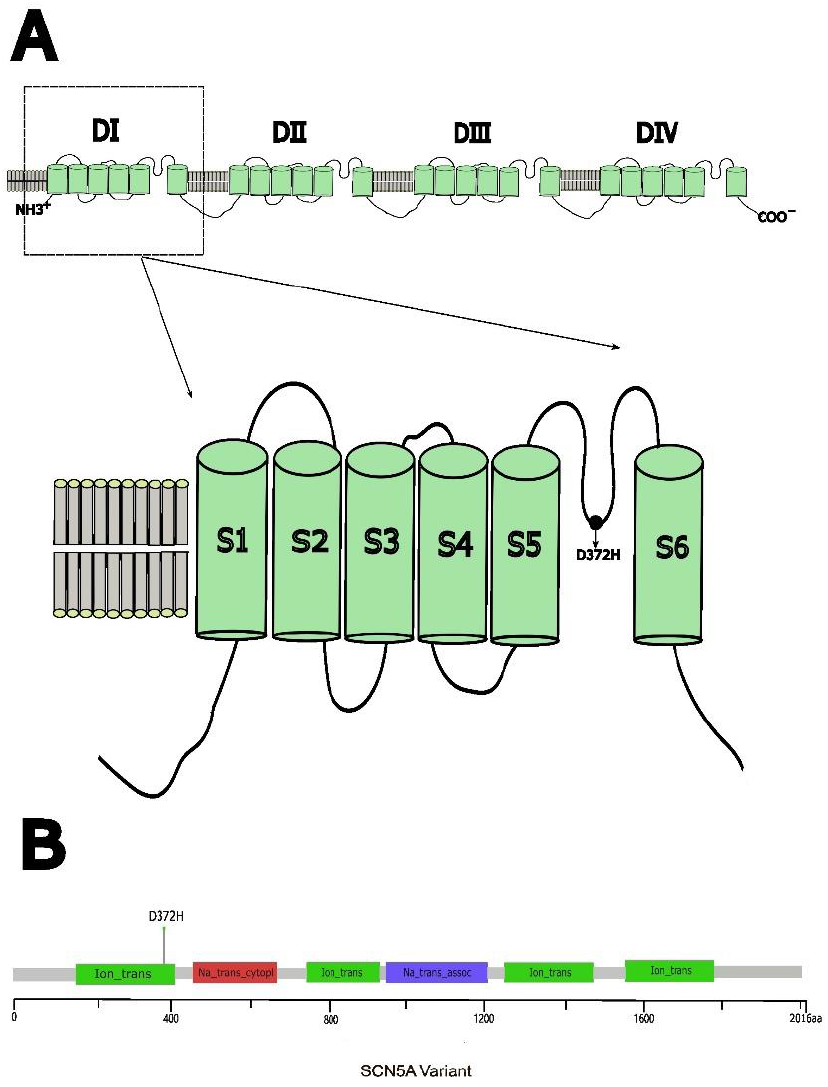
(A)Schematic drawing of the cardiac voltage-gated Na+ channel α-subunit (Nav1.5) showing the position of the D372H mutation in the linker between segments 5 and 6 in domain I (DIS5–S6). (B)The needle plot of the D372H variant in the SCN5A gene at the protein level across BrS patients.

**Fig 2.**
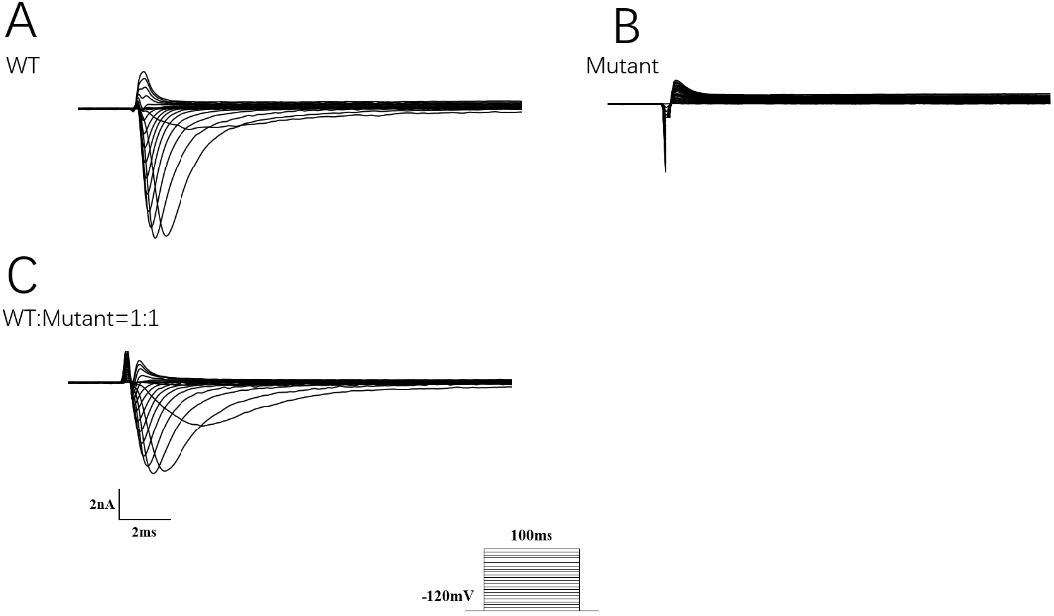
Functional Analysis of Nav1.5 Channels. (A) Current traces from HEK293 cells expressing wild-type (WT) Nav1.5 channels show robust sodium currents, indicating normal channel function with rapid activation and inactivation. (B) In contrast, traces from cells expressing the D372H mutant reveal nearly absent sodium currents, indicating a severe impairment of channel function. (C) Current traces from cells co-expressing WT and D372H mutant Nav1.5 channels at a 1:1 ratio demonstrate that despite the D372H mutation, WT channels generate significant current. I-V curve analysis shows a reduction in current density, reflecting the modulatory effects of the D372H mutation on WT channel activity.

**Fig 3.**
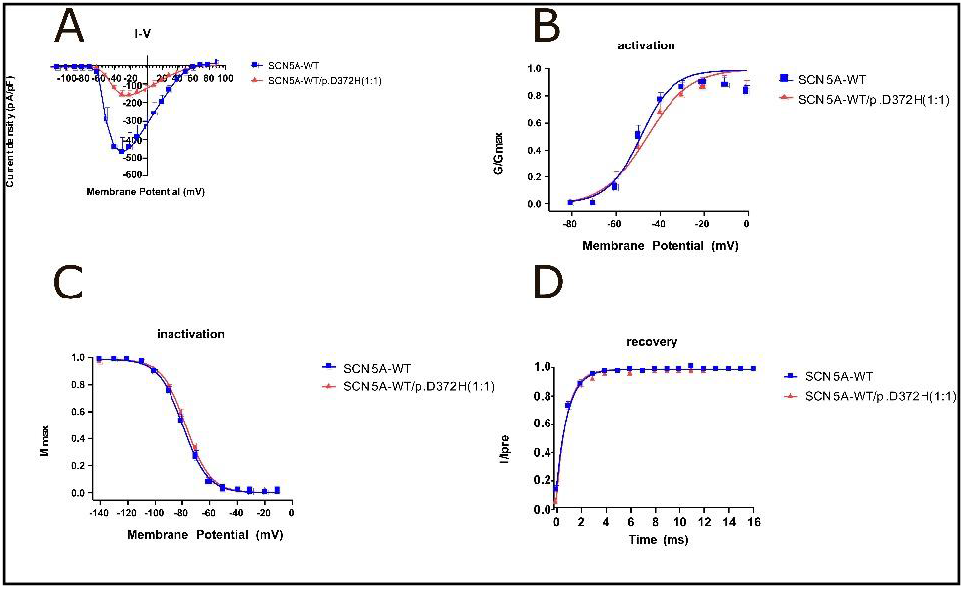
The co-expression of SCN5A-WT and SCN5A-D372H altered the steady-state activation and deactivation characteristics of sodium channels and decreased the sodium current. (A) The I-V relationship shows sodium current density in HEK293 cells expressing either SCN5A-WT (blue squares) or SCN5A-WT/p.D372H (red triangles). The curves illustrate voltage-dependent activation, with significant currents observed at depolarized potentials for both channel forms. (B) The steady-state activation curve demonstrates the normalized conductance (G/Gmax) against membrane potential. While SCN5A-WT exhibits expected activation behaviors, the D372H mutation results in a slight rightward shift, indicating altered activation properties. (C) The steady-state inactivation curve illustrates the fraction of current (I/Imax) versus membrane potential. The inactivation kinetics for SCN5A-WT and SCN5A-WT/p.D372H appear comparable, suggesting that the D372H mutation does not significantly modify inactivation behavior. (D) Recovery traces show the time-dependent recovery from inactivation, plotted as I/Ipre against recovery time. Both channel isoforms recover similarly, indicating that the D372H mutation has minimal impact on recovery kinetics.

## Materials and Methods

### 1, Plasmid constructions

According to the gene sequence of SCN5A provided by Pubmed, we designed primers on the flanks of introns to amplify the SCN5A gene by polymerase chain reaction (PCR). Mutant hNav1.5/D372H was generated using a Vazyme site-directed mutagenesis kit according to the manufacturer’s instructions. The cloning of the relevant fragment of SCN5A mutation (D372H) was generated using overlap extension PCR and inserted as an EcoR1/HindIII fragment into the EcoR1/ HindIII site of SCN5A cDNA of pcDNA3.1-EGFP-3xFLAG, which AuGCT Biotechnology provided. The head and tail primers used are:

5’-ccaagctggctagcgtttaaacttAAGCTTGCCACCATGATTCCTGGTAACCGAA-3 ‘

5’-GATGGTAGTAGAGGGATGTGGGTGCCGCTGAGAATTCtgcagatatccagcacagtggcg-3’ All constructs were purified using Qiagen columns (Qiagen Inc). The cDNA was sequenced to confirm the presence of the correct fragment, containing the mutation.

### 2, Cell culture and transient expression in HEK293 cells

HEK293 cells were grown in Dulbecco-modified Eagle’s medium (DMEM, BOSTER, Pasching, Wuhan China) supplemented with 10% fetal bovine serum (FBS, BOSTER, Pasching, Wuhan China) and 1% penicillin-streptomycin and placed at 37°C in a humidified chamber with 95% air and 5% CO2. We use HighGene transfection reagents (2μl) to transfer plasmids of pcDNA3.1-EGFP-3xFLAG (WT, 2μg) or pcDNA3.1-EGFP-3xFLAG-D372H (2μg) plasmids into cells in 6 well plates. The transfected cells were cultured at 37°C for 48 hours before performing patch-clamp, imaging, and biochemical experiments.

### 3, Electrophysiology

SCN5A-WT or SCN5A-D372H was expressed in HEK293 cells, and their channel characteristics were studied by whole-cell patch-clamp technique. I-V Curve Recording: Sodium currents were assessed by measuring current density (current per membrane capacitance) as a function of stimulation voltage. Current density was plotted on the y-axis against stimulation voltage on the x-axis.

#### Channel Characteristics

Steady-State Activation: The membrane was held at −120 mV, with test voltages applied from −80 mV to 60 mV in 10 mV increments for 100 ms. Current values were converted to conductance and normalized to the maximum conductance, generating a steady-state activation curve. The data were fitted using the Boltzmann equation Boltzmanny=1/(1+exp(-(x-x50)/-κ))to derive the half-activation voltageV_½_ and slope κ.

Steady-State Inactivation: A double-pulse protocol was used, holding at −120 mV and applying test voltages from −140 mV to 20 mV in 10 mV increments for 100 ms, followed by a 50 ms pulse at −20 mV. The steady-state inactivation curve was created by normalizing the currents to the maximum current and fitting the data with the Boltzmann equation to obtain voltage V_½_ and slope κ.

Recovery from Inactivation: A triple-pulse protocol involved holding the membrane at −120 mV, stepping to −20 mV for 100 ms, and then allowing recovery for 1 to 1000 ms before a second 50 ms pulse at −20 mV. The recovery curve was generated by normalizing the currents at different recovery times to the maximum current. Data fitting was performed using either a double-exponential function y=1 - A1exp(-x/t1) - A2exp(-x/t2) or a single-exponential function f=A1exp(-x/t1)+B, where A1 and A2 represent the proportions of fast and slow recovery components, and t1 and t2 are their respective time constants.

### 4, Confocal Laser Microscopy

To determine the subcellular localization of Nav1.5 channels, including the wild-type (WT) and the D372H mutant, in HEK293 cells, we performed confocal microscopy using Na channels conjugated to a green fluorescent protein (GFP). HEK293 cells were transfected with the plasmid containing the GFP-tagged Na channels and incubated for 48 hours. Following incubation, the cells were washed three times with phosphate-buffered saline (PBS) to remove any untransfected plasmid. The cells were then fixed in 4% formaldehyde for 30 minutes, followed by three additional washes with PBS. For nuclear staining, the slides were incubated with DAPI (BOSTER, Pasching, Wuhan, China) for 15 minutes. After incubation, the slides were carefully placed on glass coverslips using tweezers. Confocal laser microscopy (Leica) was employed to visualize the localization of the fluorescent channels.

### 5, Quantitative real-time RT-PCR analysis

To investigate the effect of D372H mutation on SCN5AmRNA expression, total RNA was isolated from cells using Trizol reagent (TaKaRa, Dalian, China), and its concentration and purity were evaluated. Using reverse transcription kits (Vazyme, Pasching, Wuhan, China) from RNA synthesis of complementary DNA (cDNA), then use q 900 and ChamQ Universal SYBR qPCR Master Mix (Vazyme, Pasching, Wuhan, China) for quantitative PCR (qPCR). The reaction mixture consisted of 2μl cDNA template, 10μl SYBR green mixture (including ROX), 200 nm forward primer and 200 nm reverse primer (total volume 20μl), and the thermal cycle conditions were: initial denaturation 30 seconds at 95°C, denaturation, annealing and extension 40 cycles—human tubulin as internal standard. The following primer sequences were listed:

SCN5A, 5’-GTCTCAGCCTTACGCACCTT-3’ (forward), 5’-GGCAGAAGACTGTGAGGACC-3’ (reverse);

TUBULIN,5’-GACAAGACCATTGGGGGAGG-3’ (forward), 5’-ACAGGCAGCAAGCCATGTAT-3’ (reverse).

The PCR products were verified by melting curve analysis, and the results were analyzed by the 2-ΔΔCt method. All experiments were performed three times.

### 6, Western Blot

To assess the expression levels of Nav1.5 channels, western blot analysis was performed on transfected HEK293 cells. The cells were seeded in six-well plates and incubated for 48 hours. Following this incubation period, IP lysate buffer was added to each well, and the plates were agitated on a shaker at 4°C for 15 minutes to extract total proteins. Protein concentrations were determined using the BCA quantification method. Subsequently, the proteins were separated using SDS-PAGE with a gradient of 4% to 12% (Boster). After electrophoresis, the proteins were transferred to a nitrocellulose membrane. The membrane was then incubated with primary antibodies against SCN5A (1:1000, Proteintech) and Tubulin (1:000, Boster) to detect the target protein and serve as a loading control, respectively. Following incubation with the primary antibodies, the membrane was washed three times with TBST (Tris-buffered saline with Tween 20) to remove unbound antibodies. The membrane was then incubated with a goat anti-rabbit secondary antibody to visualize the protein bands. For quantification, the relative expression levels of SCN5A were normalized to Tubulin, allowing for accurate comparisons of protein expression across samples.

### 7, Statistical analysis

All data are presented as means ± Standard Error of Mean (SD). Statistical comparisons between groups were conducted using appropriate tests, with significance set at p < 0.05. For experimental data, at least three independent experiments were performed, and a Student’s t-test was utilized for comparisons, with analyses conducted using SPSS version 17.0 (SPSS, Chicago, IL, USA) or GraphPad Prism software (version 9).

## Result

D372H Mutation Significantly Impairs Nav1.5 Sodium Channel Function.

To investigate the pathophysiological mechanisms underlying the phenotype of Brugada syndrome, Nav1.5 channels were transiently expressed in HEK 293 cells, and the functional impact of the D372H mutation was assessed using whole-cell patch-clamp recordings. Initial results indicated that sodium currents in cells transfected with the D372H mutant nearly disappeared. To further explore the effects of this mutation, we designed a subsequent patch-clamp experiment where equal amounts of wild-type (WT) and D372H plasmids were co-transfected into HEK293 cells at a 1:1 ratio. Patch-clamp recordings were then performed specifically on cells predominantly expressing the WT channels. This strategy enabled us to evaluate the functional properties of the WT channels in the presence of the D372H mutant, providing insights into potential interactions or modulatory effects of the mutant on overall current. Analysis of the currents recorded from WT-expressing cells revealed a significant reduction in current density after co-expression with the D372H mutation, as demonstrated by the I-V curve. This finding indicates that the SCN5A-WT/p.D372H (1:1) co-expression leads to the down-regulation of channel activity, with statistical significance observed between the groups (P < 0.05).

Table 1 summarizes the current density measurements of sodium channels at −20 mV for two groups: SCN5A-WT and SCN5A-WT/p.D372H (1:1). The SCN5A-WT group (n=8) exhibited a current density of −473.10 ± 82.27 pA/pF, whereas the SCN5A-WT/p.D372H group (n=8) displayed a significantly lower current density of −158.54 ± 28.37 pA/pF (p < 0.05). This reduction in current density in the D372H mutant indicates a substantial impairment in sodium channel function compared to the wild-type. The I-V curve depicted in the accompanying figure provides a visual representation of the voltage-dependent properties of these channels. The SCN5A-WT group, indicated by the blue line, shows a robust inward sodium current with a pronounced peak, while the red line representing the D372H mutant demonstrates markedly diminished current amplitudes. This graphical data reinforces the statistical significance noted in the table, highlighting that the D372H mutation significantly reduces sodium current. The activation characteristics of sodium channels were compared between SCN5A-WT and SCN5A-WT/p.D372H (1:1) groups as shown in Table 2 and the corresponding activation curve. The half-activation voltage (V1/2) for SCN5A-WT was measured at −47.16 ± 2.91 mV, whereas the D372H mutant showed a slightly more depolarized V1/2 at −45.50 ± 4.21 mV. This suggests that the D372H mutation shifts the voltage dependency of activation toward more positive potentials, indicating that more depolarization is required to activate the sodium channels effectively. The slope factor (κ) for SCN5A-WT was 8.06 ± 2.33 mV, reflecting a steep activation curve, compared to a lower slope of 6.36 ± 1.65 mV for the D372H mutant. A lower slope indicates that the D372H mutation results in less efficient activation over a range of voltages. However, the surface D372H mutation did not significantly change sodium channels’ inactivation dynamics and recovery dynamics.

**Table 1.**
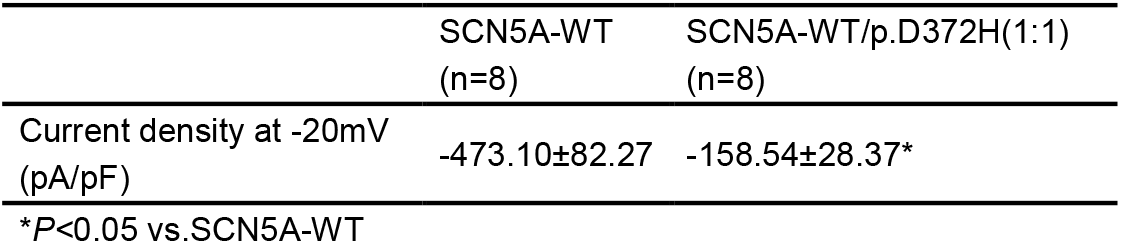
I-V.

**Table 2.** activation.

| Table 2. activation |  |  |
| --- | --- | --- |
|  | SCN5A-WT<br>(n=8) | SCN5A-WT/p.D372H(1:1)<br>(n=8) |
| V1/2 (mV) | -47.16±2.91 | -45.50±4.21 |
| $\kappa$ (mV) | 8.06±2.33 | 6.36±1.65 |

### Impaired Membrane Localization of NaV1.5 in SCN5A-D372H Mutants

To further investigate the effect of the SCN5A-D372H mutant on the functional expression of the NaV1.5 sodium channel protein, we employed confocal microscopy to observe the transfected HEK293 cells. Figure 4A illustrates that SCN5A-WT predominantly localizes to the cell membrane, exhibiting a strong green fluorescent signal. In contrast, the SCN5A-D372H mutant displayed markedly weaker fluorescence at the cell membrane, with the protein exhibiting a relatively dispersed distribution throughout the cytoplasm.

**Fig 4.**
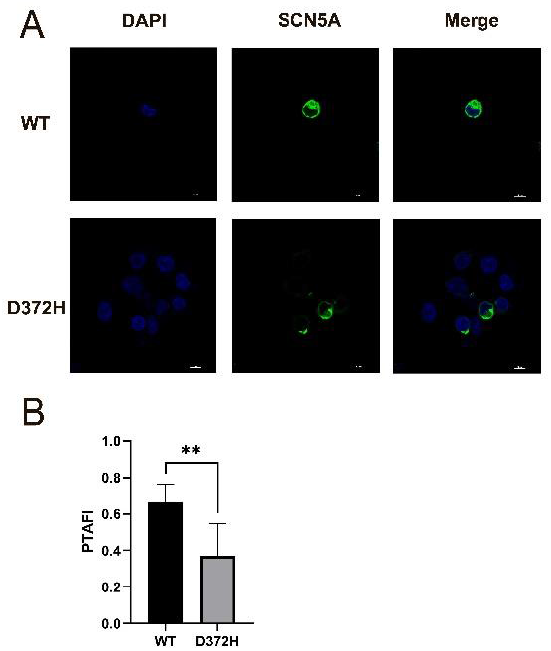
(A) A representative confocal microscope scan showed the localization of Na channels in 293T cells. The WT channels conjugated with GFP were stained both outside and in the centre of the cell, suggesting that the WT channels were generated in the centre of the cell and migrated to the cell membrane. Cells transfected with D372H mutant plasmid showed weaker fluorescence on the cell membrane than WT, and the protein distribution in the cytoplasm was relatively dispersed. (B) Comparison of PTAFI (the ratio of membrane fluorescence to total cellular fluorescence) between wild-type (WT) and D372H mutant groups. Bars represent mean values with error bars indicating the standard error of the mean. n=6 in each group. ∗ ∗P< 0.01, mean SD.

To quantitatively assess these observations, we calculated the PTAFI (the ratio of membrane fluorescence to total cellular fluorescence) for both wild-type (WT) and D372H mutant groups. The results indicate that the WT group has a significantly higher PTAFI (mean ± SD) compared to the D372H mutant group, suggesting that the D372H mutation adversely affects the membrane localization of the protein (p < 0.05) (Figure 4B).

### D372H Mutation Reduces Nav1.5 mRNA and Protein Expression Levels

The impact of the D372H mutation on Nav1.5 expression was investigated at the mRNA level using RT-qPCR. The results demonstrate that the WT group exhibits significantly higher Nav1.5 mRNA levels compared to the D372H mutant (p < 0.001). This indicates that the D372H mutation results in a substantial reduction in Nav1.5 transcript levels, which may impair its functional role in cellular processes(Fig.5A). To further investigate the expression of Nav1.5 at the protein level, Western blot assays were conducted. The results revealed that the total cell expression of the D372H mutant Nav1.5 channels was markedly lower than that of the WT channels, reinforcing the findings from the mRNA analysis. This suggests that the D372H mutation not only affects transcript levels but also leads to reduced protein expression at the plasma membrane(Fig.5B).

**Fig 5.**
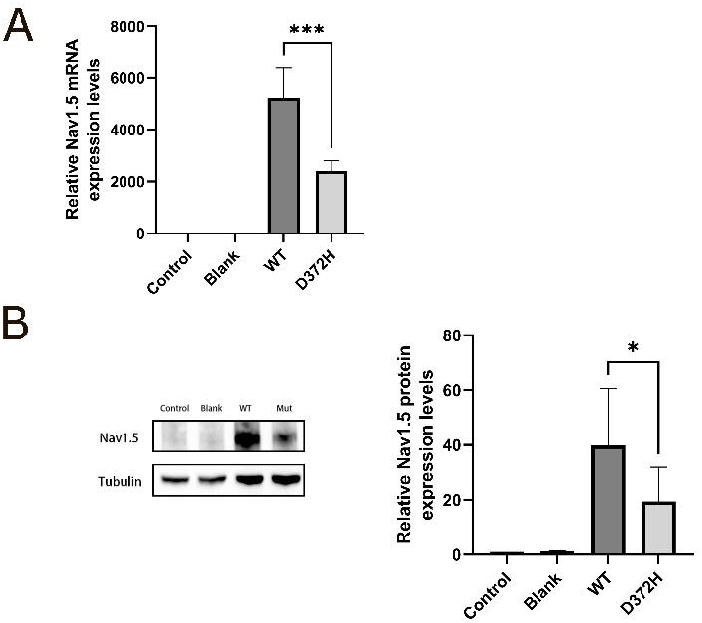
Effects of the D372H mutation on Nav1.5 expression levels. (A) Relative Nav1.5 mRNA expression levels measured by RT-qPCR. The data indicate that the WT group exhibits significantly higher Nav1.5 mRNA levels compared to the D372H mutant (^∗∗∗^p < 0.001). The Control group shows mRNA expression comparable to the WT. (B) Western blot analysis of total Nav1.5 protein expression. The left panel displays representative blots for Nav1.5 and the loading control, Tubulin. The right panel quantifies relative Nav1.5 protein expression levels. The WT group shows higher protein expression than the D372H mutant. Data are presented as mean ± SD. ^∗^P < 0.05

## Discussion

This study provides a comprehensive analysis of the functional and molecular effects of the D372H mutation in the Nav1.5 sodium channel, which is associated with Brugada syndrome. Through a combination of electrophysiological recordings, confocal microscopy, and molecular assays, we elucidated how the D372H mutation impacts channel activity, expression, and localization. Our electrophysiological studies using whole-cell patch-clamp recordings demonstrated that sodium currents in HEK293 cells expressing the D372H mutant were nearly absent. The co-expression experiments, where equal amounts of WT and D372H plasmids were transfected, revealed that in the presence of the D372H mutation, the current density was markedly reduced, suggesting that the D372H mutation exerts a dominant-negative effect on WT channel function. This reduction in current density was statistically significant, highlighting the modulatory influence of the D372H mutation on sodium channel activity. Confocal microscopy provided insights into the localization of Nav1.5 channels. The strong membrane localization of WT channels contrasted sharply with the weaker fluorescence and more dispersed cytoplasmic distribution of the D372H mutant. At the molecular level, our RT-qPCR and Western blot analyses revealed a significant reduction in both Nav1.5 mRNA and protein levels in cells expressing the D372H mutant compared to WT. This suggests that the D372H mutation not only disrupts channel function but also affects its expression at the transcriptional and translational levels. The marked decrease in transcript levels indicates potential disruptions in transcriptional regulation or mRNA stability, while the reduced protein expression at the plasma membrane reinforces the notion that the D372H mutation leads to impaired functional expression of Nav1.5.

The SCN5A gene is a critical player in the pathophysiology of Brugada syndrome (BrS), with over 300 mutation sites identified^19^. However, the majority of these mutations are primarily documented through clinical reports, lacking robust in vitro functional experiments to substantiate their effects. Previous studies have investigated several mechanisms by which missense mutations can lead to reduced sodium current density, identifying a range of contributing factors^18^. Certain mutations can result in diminished expression levels of the Nav1.5 sodium channel. This reduction can significantly impact the overall sodium current available for cardiac action potentials, thereby affecting the excitability and conduction properties of cardiac tissue. Mutations may disrupt the normal processes of activation and inactivation of sodium channels. Such disruptions can compromise the channel’s ability to open and close in response to voltage changes, ultimately impairing its physiological function and contributing to arrhythmogenic events. Some mutations may induce conformational alterations within the channel structure, leading to a loss of functional capacity. These structural changes can hinder the channel’s ability to effectively conduct sodium ions, thereby increasing the risk of arrhythmias. Mutations can interfere with the transport of the Nav1.5 channel from the endoplasmic reticulum (ER) and Golgi apparatus to the plasma membrane. For example, specific mutations such as R1432G^20^ and D1690N^21^ have been shown to result in improper transport of the mutant channels, causing their retention in the ER due to cellular quality control mechanisms.

### Limitation

The results of this study indicate that the SCN5A-D372H mutation, located in a critical domain of the NaV1.5 sodium channel, leads to functional impairment characterized by reduced sodium current density and decreased channel expression levels, potentially contributing to the pathogenesis of Brugada syndrome. However, several limitations should be noted. Firstly, while we observed a decrease in sodium current density associated with the D372H mutation, the specific mechanisms underlying this loss remain unclear, necessitating further research to elucidate the affected molecular pathways. Secondly, we utilized HEK293 cells for our experiments, which, although useful for studying ion channel properties through plasmid-mediated overexpression, do not fully replicate the physiological environment of cardiac myocytes. As a result, factors such as polygenic modifiers relevant to Brugada syndrome may not be adequately represented, limiting the applicability of our findings to native cardiac tissues^22^. Lastly, earlier research has demonstrated that the W374G mutation results in a decrease in sodium current density, which can be partially recovered with mexiletine (MEX)^18^. This suggests that trafficking defects associated with SCN5A mutations in the pore-S6 regions of Domains I, III, and IV can be rescued by MEX, whereas mutations in Domain II do not exhibit this property. We hypothesize that the current loss associated with the D372H mutation may also be amenable to rescue by MEX, warranting further investigation into its therapeutic potential in Brugada syndrome.

## Conclusion

In conclusion, our study highlights the multifaceted impact of the D372H mutation on Nav1.5 sodium channels, providing a clearer understanding of its role in the pathogenesis of Brugada syndrome. These findings underscore the importance of further research to explore therapeutic strategies that could mitigate the functional deficits associated with this and similar mutations.

## Supplemental figure

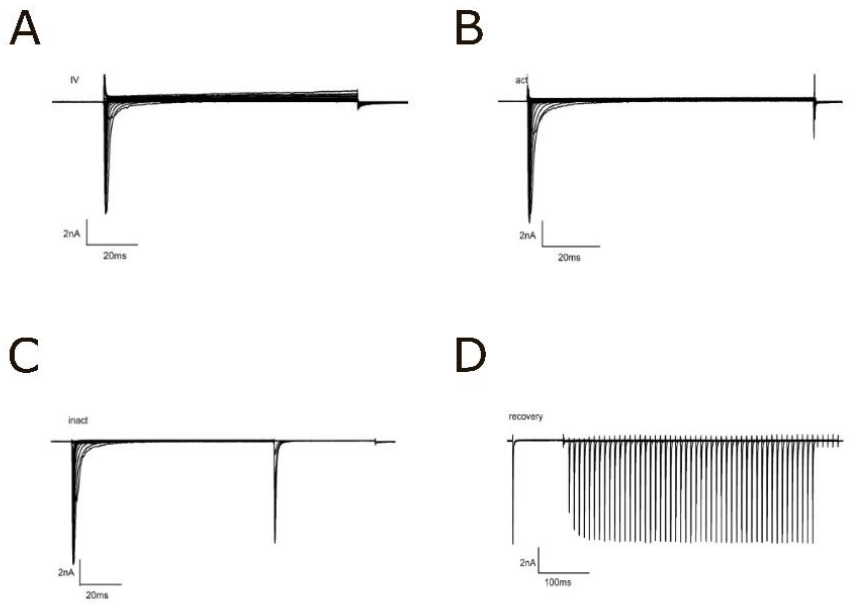

This diagram shows the electrophysiological characteristics of sodium channels under normal conditions, and the typical current diagram of sodium channel activation, inactivation, and recovery after inactivation is recorded by changing the voltage.

